# D-lactate derived from intestinal bacteria drives lysine D-lactylation to modulate transcription in liver cells

**DOI:** 10.1101/2024.10.04.616636

**Authors:** Yong Zang, Jianji Zhang, Mingxin Xia, Aiyuan Wang, Zilong Fan, Yanpu Han, Hui Zhang, Siyu Wang, Ziping Niu, Jingya Wu, Xue Bai, Shanshan Tian, Chen Chen, Guijin Zhai, Hanyang Dong, Deqing Hu, Zhongyi Cheng, Kai Zhang

## Abstract

Emerging evidence indicates that the accumulated D-lactate (D-la) from intestinal bacteria induces diverse effects in human intestinal injury diseases. However, its regulatory mechanism and functional consequence remain unclear. Here, we show that the secreted D-la induces lysine D-lactylation (K-Dla) to modulate transcription in host cells. We uncover K-Dla in HepG2 cells stimulated by D-la from *E. coli* secretions. Then, we confirm the occurrence of D-la-driven K-Dla by four orthogonal approaches. We further validate that the existence of K-Dla in organisms by detecting the mouse liver and human intestinal tissue. Besides, we demonstrate a SCOT1-catalyzed biosynthetic pathway of D-lactyl-CoA, serving as a key donor for K-Dla. Next, we identify the K-Dla landscape with 2895 sites and further suggest the regulation of histone K-Dla in transcription by MNase ChIP-seq and RNA-seq assays. Finally, we show the potential effect of K-Dla on inhibiting the hepatocellular carcinoma cell proliferation. In summary, we report that the secreted D-la can induce K-Dla to regulate transcription, providing a new insight into the function of D-la secreted by gut microbiota and the effect of protein chemical modifications induced by bacterial secretions.

## Introduction

Increasing evidence suggests that abnormal metabolites can diver intricately disease progression.^1, 2, 3^ As an important metabolite in the human body, lactate exists as two optical enantiomers: L-lactate (L-la) and D-lactate (D-la), however, they have fundamental differences in terms of source, metabolic pathway, as well as chemical and biological effects. Distinguished from L-la, D-la is typically present in a relatively low content in the human plasma, primarily deriving from intestinal bacterial metabolism and food consumption or endogenously methylglyoxal metabolism pathway.^4^ However, in certain human intestinal injury diseases such as short bowel syndrome and Crohn’s disease, larges amount of D-la can is produced and secreted by intestinal bacteria such as *E. coli* in the hypoxic intestinal microenvironment, then enter human intestinal cells and circulate through the bloodstream to liver.^5, 6^ The secreted D-la can accumulated to the levels of 7 mM or higher in the human body,^7^ and thereby inducing some effects on digestive system^8^, even affecting neurological symptoms and immunosuppressive tumor microenvironment (TME).^7, 9, 10^ However, the regulatory mechanism and functional consequence of this secreted D-la in host cells remains largely unclear.

As the enantiomer of D-la, L-la has recently been rediscovered as an important signaling molecule, serving both metabolic and non-metabolic functions.^11, 12, 13, 14^ The recently identified histone lysine L-lactylation (K-Lla) induced by L-la has been believed play a key regulatory role in tumor microenvironment, providing a new insight into the non-metabolic functions of L-la.^15^ Meanwhile, recent study shows that D-lactylation (K-Dla) may occur via a non-enzymatic acyl transfer from S-D-(*R*)-lactoylglutathione, deviating from the glycolytic by-product, to regulate the metabolic output.^16^ However, it remains unclear whether the secreted D-la induces the generation of K-Dla for regulation of host cells.

Herein, we propose a hypothesis that D-la secreted by the gut microbiota can accumulate in liver in intestinal injury diseases and induce lysine D-lactylation (K-Dla) to regulate transcription in host. We first discovered K-Dla in HepG2 cells stimulated by D-la from *E. coli* secretions in a hypoxic environment. We then confirmed the occurrence of K-Dla induced by D-la using four orthogonal approaches, including high performed liquid chromatography (HPLC) retention time, MS/MS fragmentation pattern, immunoblot and immunofluorescence assays based on pan anti-K-Dla antibody, as well as metabolic labelling experiments using isotopic sodium D-lactate (^13^C1-D-la). We further validated that the existence of K-Dla in organisms by detecting the mouse liver and human intestinal tissue. Next, we revealed a D-la-driven K-Dla pathway, in which SCOT1 can catalyze the transformation of D-la to D-lactyl-CoA, serving as a key donor for generation of K-Dla. Furthermore, we identified the K-Dla landscape with 2895 sites using the K-Dla enrichment coupled with LC-MS/MS analysis and characterized the K-Dla localization, enriched molecular network and pathways. We showed that D-la-driven K-Dla is a unique post-translational modification, distinguished from K-Lla and K-Dla from S-D-(*R*)- lactoylglutathione. Besides, we identified 34 K-Dla sites on core histones obviously activated by D-la using a SILAC based quantitative proteomics approach. MNase ChIP-seq, and RNA-seq assays further indicated the function of K-Dla in transcription regulation.

Finally, we explored the effect of K-Dla on inhibiting the proliferation of hepatocellular carcinoma cells. In summary, we uncover the secreted D-la-driven K-Dla, elucidate its formation mechanism and landscape, and reveal a transcription regulation mediated by K-Dla for tumor suppression, providing new insight into the protein chemical modifications induced by gut microbiota secretions.

## Results

### Identification and validation of D-lactate-induced lysine D-lactylation in cells

D-lactate (D-la) is typically present in the adult blood at nanomolar concentrations.^17^ However, in certain diseases such as short bowel syndrome, the concentration of D-la may rapidly rise to millimolar levels due to the entry of D-lactate secreted by intestinal microflora into human cells.^18, 19^ Although the effects of D-la are recognized, its regulatory mechanism remains unclear. Here, we hypothesized that the secreted D-la acts as a precursor for lysine D-lactylation (K-Dla) that regulates the fates of cells by remodeling the local chromatin environment.

To simulate the intestinal hypoxia environment, we cultured Escherichia coli (*E. coli*) *MG 1655* in anaerobic medium, and thus detected the millimolar concentration of D-La in the supernatant of *E. coli* (Fig. 1b), while L-la was nearly undetectable (Fig. 1c). To explore whether the secreted D-la drives generation of K-Dla, we used D-la to stimulate the HepG2 cells and performed a mass spectrometry-based proteomic analysis. As a result, we observed a mass shift of 72.021 Da at lysine residues in a proteolytic peptide, which was absent in non-stimulated samples (Fig. 1d). The mass shift falls into the molecular weight of K-Dla, suggesting that D-la may induce the addition of D-lactyl group to the ε-amino group of lysine residue.

**Fig. 1.**
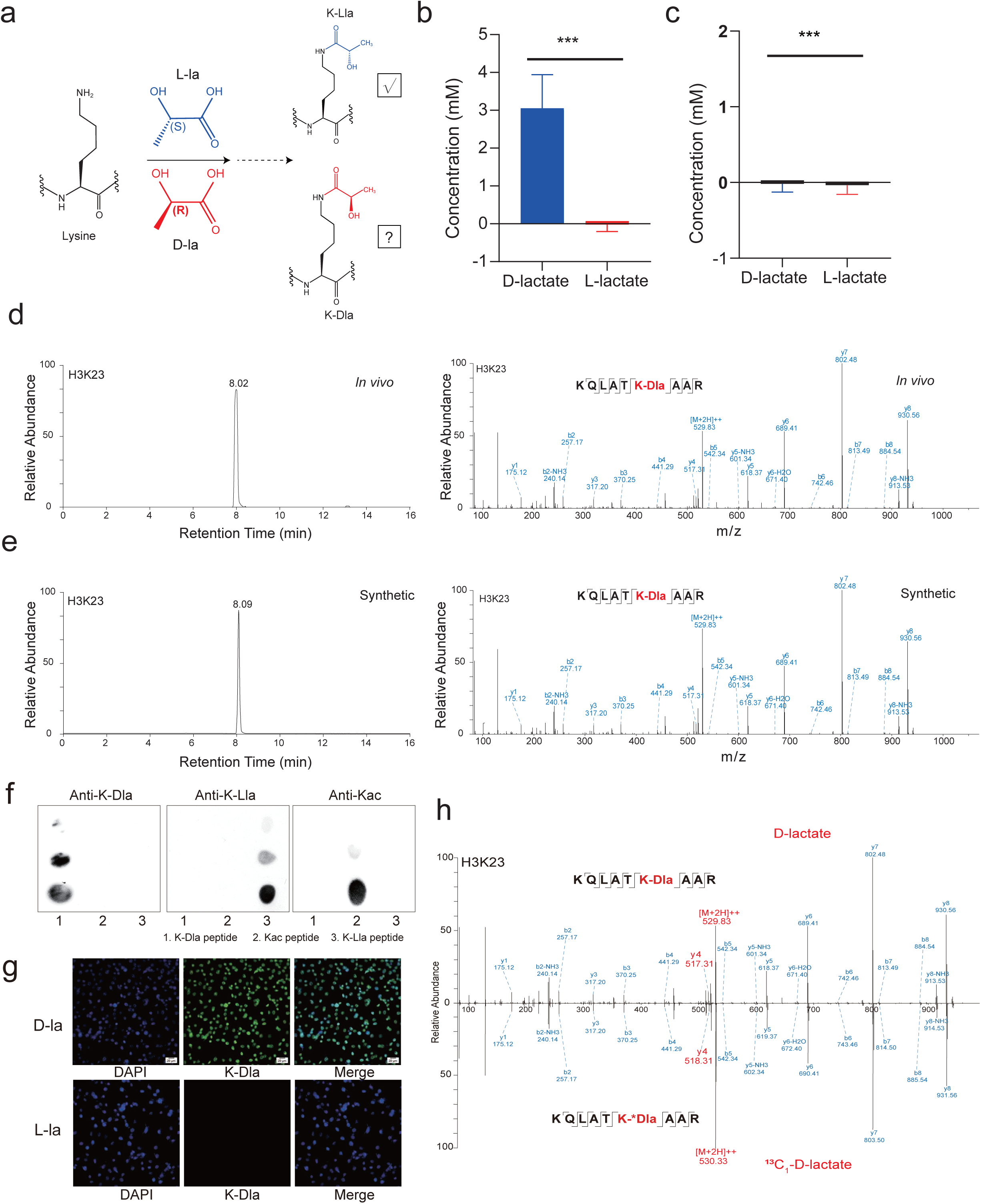
The identification and validation of D-lactate-driven lysine D-lactylation in cells. a, Chemical structure of K-Dla and K-Lla. b, The concentration of D-lactate secreted by E. coli MG 1655 in hypoxia environment. c, The concentration of L-lactate secreted by E. coli MG 1655 in hypoxia environment. d, e, f, MS/MS spectra of a lactylated histone peptide (H3K23 K-Dla) derived from Hepg2 cells which were stimulated by D-la (in vivo), its synthetic counterpart and its H3K23ac counterpart (in vivo). g, Antibody specificity was analyzed by dot blot assay. i, Immunofluorescence microscopy images of cells treated with D-la or L-la. Blue: DAPI; Green: K-Dla. Hela cells. h, MS/MS spectra of light (upper) and heavy (lower) K-Dla peptide (H3K23). The ^13^C1-K-Dla peptide was identified in HepG2 cells treated with 10 mm ^13^C1-D-la.

To validate the D-la-induced K-Dla, we carried out a set of biochemical assays. First, we chemically synthesized the K-Dla peptide (histone H3K23 K-Dla, KQLATK (D-la) AAR), and compared its MS/MS fragmentation and LC retention time with its in vivo-derived counterpart. The synthetic peptides showed nearly chromatographic retention time and MS/MS fragmentation pattern to the corresponding endogenous peptide (Fig. 1d, e and Extended Data Fig1. a).

Next, we generated a pan antibody against D-lactyllysine to confirm the specificity of K-Dla. The results of dot-blotting assays showed that pan anti-D-lactyllysine antibody can specifically bind with the K-Dla peptide, while K-Dla peptide can’t bind with the other types of pan lysine modification antibody (Fig. 1f), demonstrating the specificity of the pan anti K-Dla antibody. Subsequently, we used immunofluorescence assays to detect HepG2 and Hela cells, which were stimulated by D-la or L-la, respectively. The results showed that only D-la stimulated cells can be observed to exhibit K-Dla fluorescence by confocal microscopy (Fig. 1g and Extended Data Fig. 1a), supporting that the K-Dla is indeed induced by D-la. Notably, we found that K-Dla fluorescence is highly focused in the nucleus, suggesting that K-Dla substrates proteins are enriched in nucleus.

To confirm further that the secreted D-la acts as the precursor for K-Dla production, we performed a metabolic labelling experiment using isotopic labeling D-la (^13^C1-D-la) followed by MS/MS analysis to distinguish the exogenous D-la, L-la and endogenous D-la. Sodium D-lactate (^13^C1) was exerted to stimulate HepG2 cells for 48 h, then the proteins were digested by trypsin and enriched with anti-D-lactyllysine antibody-agarose beads followed by LC-MS/MS analysis. We identified K-Dla peptides bearing ^13^C1-D-la, including the H3K23 K-Dla peptide (Fig. 1h), convincingly demonstrating that D-la acts as a metabolic precursor for K-Dla.

Taken together, these results demonstrate that the secreted D-la can induce the generation of K-Dla in host cells.

### The molecular mechanism by which D-la induces lysine D-lactylation

The D-la mainly was secreted by gut microbiota in human body, so we used immunohistochemical to test the level of K-Dla in human colon tissue. The results showed that D-Kla proteins were concentrated in the villus layer (Fig. 2a). It has been known that oxygen concentrations along the mammalian intestinal tract are lower than in other tissues,^20^ and intestinal absorption of D-la may be stored or metabolized through the liver.^6^ Thus, we speculate that bacteria in the gut environment could secrete D-la to influence the abundance of D-Kla in liver. So, we used the anaerobic medium from *E. coli MG 1655* to stimulate the HepG2 cells and found the hypoxia conditioned medium increased the level of K-Dla (Fig. 2b and Extended Data Fig.2a). Furthermore, we used antibiotics to clear the gut microbiota in mice and found the level of D-Kla were decreased accordingly in liver (Fig. 2c and Extended Data Fig. 2b). These data indicate that the D-la secreted by gut microbiota can drive the generation of K-Dla in the host’s liver and intestinal tissues.

**Fig. 2.**
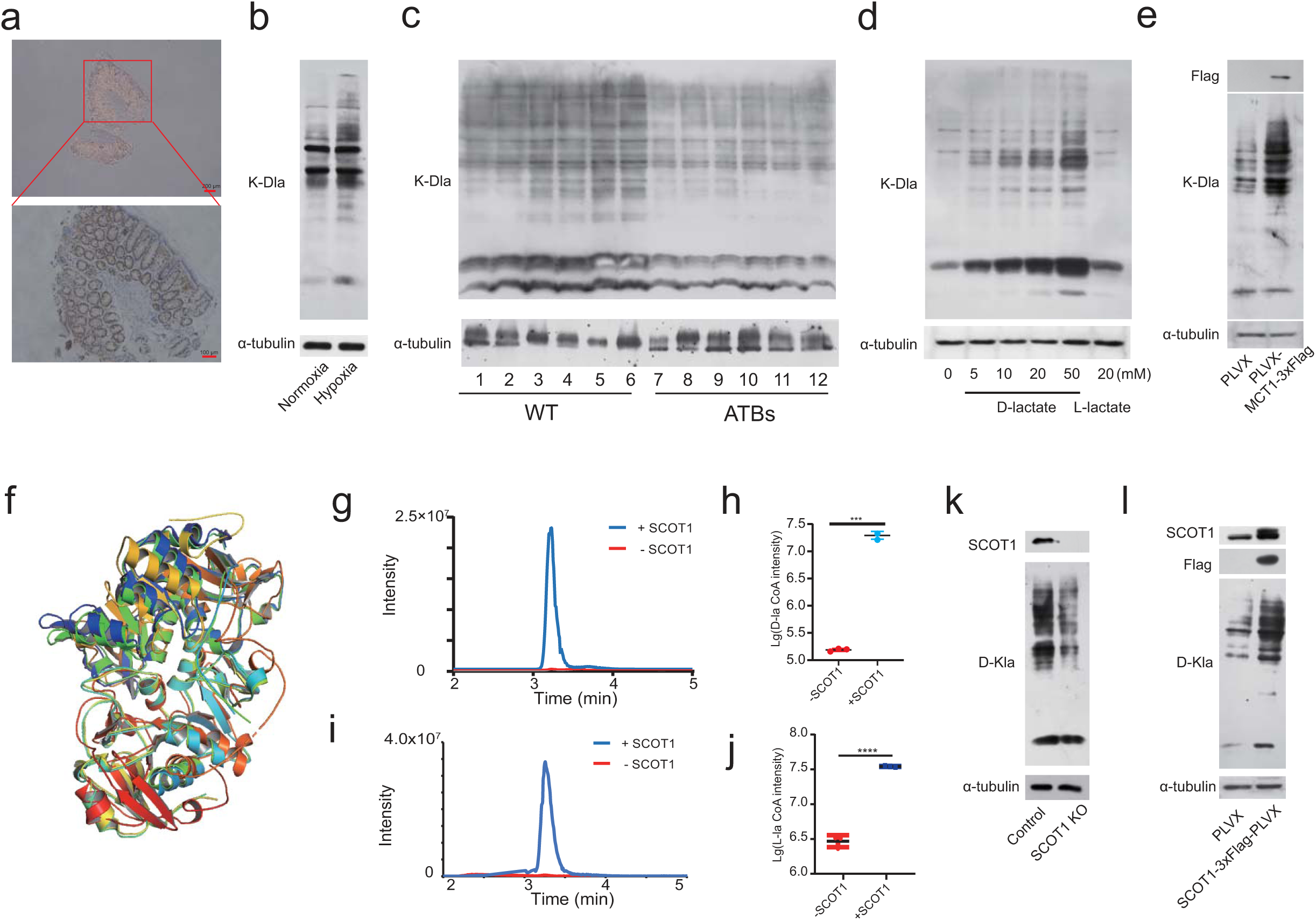
The progression of D-la to lysine D-lactylation. a, Representative IHC of K-Dla in colon tissue. b, The level of K-Dla in cells which were stimulated by anaerobic medium from *E. coli MG 1655* cultured in normoxia or hypoxia environment. c, The level of K-Dla in mice liver (the gut microbiota were cleared up by antibiotics (ATBs) or not. d, The level of K-Dla in cells which were cultured with indicated doses of D-la or L-la for 48h. e, The level of K-Dla in overexpressed MCT1 cells which were cultured with 5 mm D-la for 48h. f, Compare the structures of human SCOT1, pig heart SCOT1 and YdiF. g, h, i, j is performed produce D/L-lactyl-CoA (D-la-CoA) by SCOT1. − SCOT1: Incubation D/L-lactate with succinyl-CoA, + SCOT1: incubation D/L-lactate with succinyl-CoA with SCOT1. n=3 biological repetitions. k, l, The level of K-Dla in overexpressed or knockout SCOT1 cells which were cultured with 5 mm D-La for 48h.

We next aimed to explore how D-la is converted to K-Dla. We stimulated the HepG2 cells with sodium D-lactate and sodium L-lactate, respectively. Western blot assays showed that D-la increased the abundance of K-Dla, slightly decreased K-Lla level and did not significantly affect the level of Kac (Fig. 2b and Extended Data Fig. 2b, c), suggesting the K-Dla can be induced by the secreted D-lactate from *E. coli* in hypoxic environment.

To determine whether the intracellular K-Dla is stimulated by external D-lactate, we investigated the transmission mechanism of D-la. It is known that both of D-la and L-la can be transported across the plasma membrane through MCT1 and MCT4, with MCT1 being responsible for D-la uptake into cells.^21, 22, 23^ Compared to wild type HepG2 cells, we found that K-Dla level was obviously increased in HepG2 cells overexpressing MCT1, suggesting that K-Dla is induced by external D-la (Fig. 2c and Extended Data Fig. 2d).

We further set out to investigate whether the D-la can be converted chemically to D-lactyl-coenzyme A, a potential donor for K-Dla. Our recent study showed that YdiF, a CoA transferase in *E. coli*, can catalyze the formation of a lactyl-coenzyme A. Inspired by it, we used SHF(https://micc.tmu.edu.cn/shf) to analogue protein structure between YdiF and human proteins, and found that SCOT1 has a 75.3% similarity with YdiF. To observe the structure in the microstructure, we also used the PyMol to compare the human SCOT1, pig heart SCOT1 and YdiF, revealing their high structural similarity (Fig. 2d). We therefore reasonably speculate that SCOT1 has a potential catalytic activity for transferring D-la to D-lactyl-coenzyme A. We incubated purified SCOT1 with sodium D-lactate and sodium L-lactate, respectively, and observed that SCOT1 could catalytically transfer CoA group from a succinyl-CoA to the D-la or L-la and thus produce the D-lactyl-CoA or L-lactyl-CoA in vitro (Fig. 2e-h and Extended Data Fig. 2e), indicating that SCOT1 has a lactyl-CoA-transferase activity. To confirm SCOT1 has intracellular catalytic activity, we further tested the K-Dla and K-Lla levels in cells overexpressed or knockout (KO) SCOT1. Compared to the control group, overexpressed SCOT1 caused obviously increases in K-Dla and K-Lla levels, while KO SCOT1 significantly decreased K-Dla and K-Lla levels (Fig. 2i, j and Extended Data Fig. 2f, g). These results reveal the biosynthetic pathway from D-la to D-lactyl-CoA.

Together, our results showed that the secreted D-lactate can be transported into cells by MCT1 and converted chemically to D-lactyl-CoA under the catalysis of SCOT1, which acts as a key step for final produce of K-Dla on protein substrates.

### Systematic profiling of the K-Dla in HepG2 cells

Distinguished from the pathway of lactoylglutathione, the substrates of D-la-induced K-Dla and its functional consequence remain unknown. To this purpose, we performed a proteomic analysis by combining K-Dla affinity enrichment with mass spectrometry identification for the D-lactylome induced by exogenous D-la in HepG2 cells. Surprisingly, a landscape of K-Dla was presented with 2895 high-confidence K-Dla sites on 1215 proteins (Fig. 3a, Extended Data Fig. 3a, b and table S1). To better characterize the protein modification, we analyzed the subcellular localization of the identified K-Dla substrates proteins using the wolfpsort. Interestingly, the D-lactylome analysis showed that K-Dla substrates are widespread in cells, with 59.2% of proteins localized to the cell nucleus, consistent with our immunofluorescence observation (Fig. 3c, d). To explore the biological significance of K-Dla substrates proteins, we further performed Gene Ontology (GO) analysis, suggesting that K-Dla substrates proteins are mainly enriched in RNA related biological processes, especially mRNA progressing and RNA splicing (Extended Data Fig. 3d). This finding was further supported by biochemical pathway analysis of Kyoto Encyclopedia of Genes and Genomes (KEGG) and Reactome pathways enrichment analysis (Extended Data Fig. 3c, e).

**Fig. 3.**
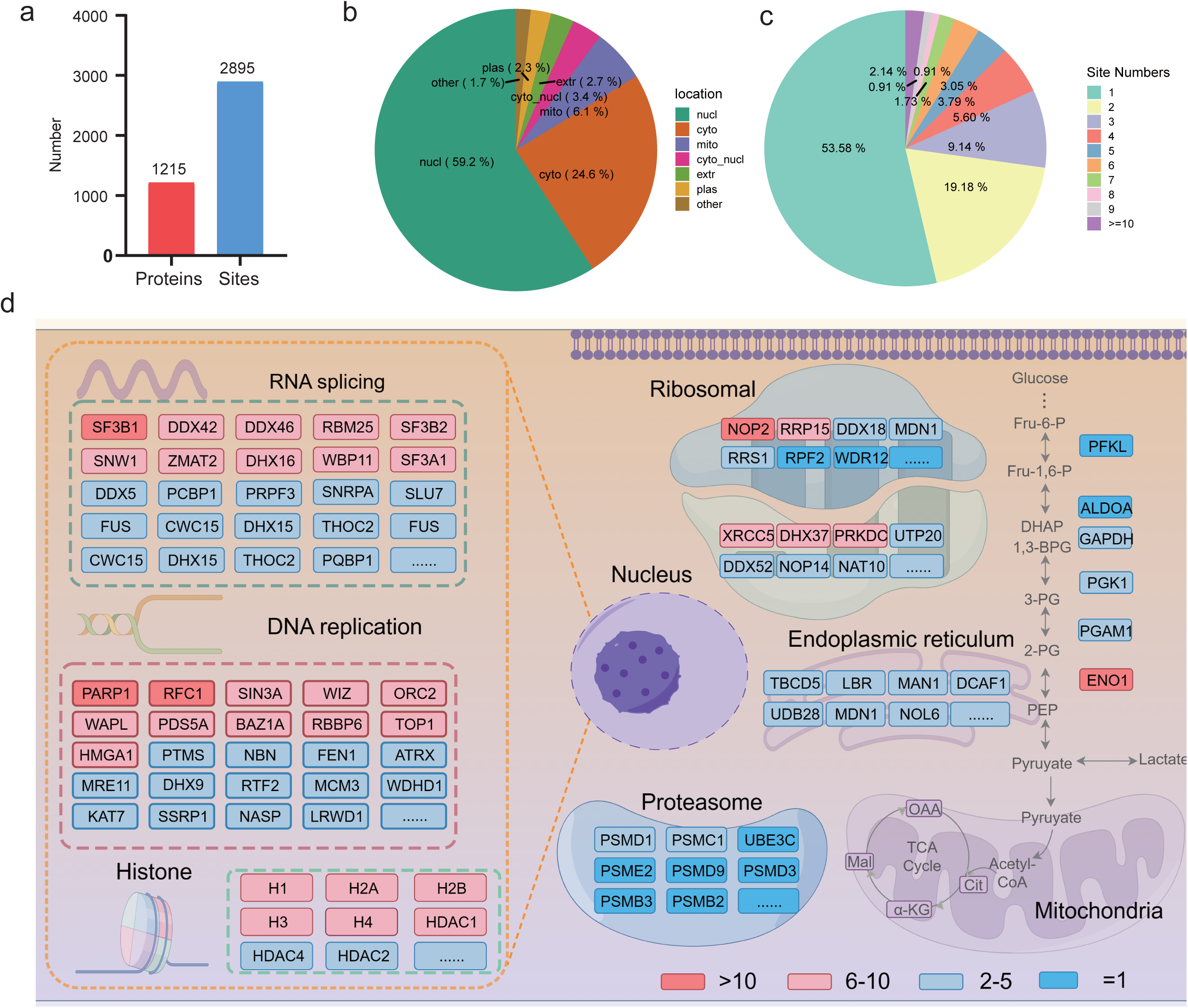
Characterization of the K-Dla proteome in HepG2 cells. a, Statistics analysis of the K-Dla proteins and sites of HepG2 cells. b, Subcellular localization of the K-Dla proteins in HepG2 cells. c, The frequency of K-Dla occurs on proteins in HepG2 cells. d, Overview of K-Dla proteins in cytoplasmic organoids based on proteome and D-lactylome in HepG2 cells.

To further characterize the D-lactylome, we performed a systematic comparison with L-lactylome from HCC patients.^24^ Interestingly, we found that there is a significant difference in the distribution of their substrate proteins. Different from D-la substrates in nucleus, K-Lla substrates are mainly located in the cytosol and enriched in metabolism pathways, such as amino acid metabolism, carbon metabolism and fatty acid metabolism (Extended Data Fig. 3f-j). The results indicate that K-Dla is likely to have unique functions and chemical biology effects distinguished from K-Lla.

### Identification of histone K-Dla induced by D-la

Given that histone modifications typically play key roles in multiple processes and K-Dla is enriched in nucleus, we next examined the histone K-Dla in nucleus. We performed a SILAC-based quantitative proteomics approach to characterize the levels of K-Dla on histone proteins in HepG2 cells stimulated with D-la or not (Fig.4a). As a result, 61 K-Dla peptides were identified and quantified on histone proteins in D-la group including 8 sites on H3, 5 sites on H4, 4 sites on H2A and 13 sites on H2B, whereas the abundance of K-Dla in control groups was very low (Fig. 4b, c and table S2), suggesting the D-la may be transferred to nucleus and drive the generation of K-Dla on histones.

**Fig. 4.**
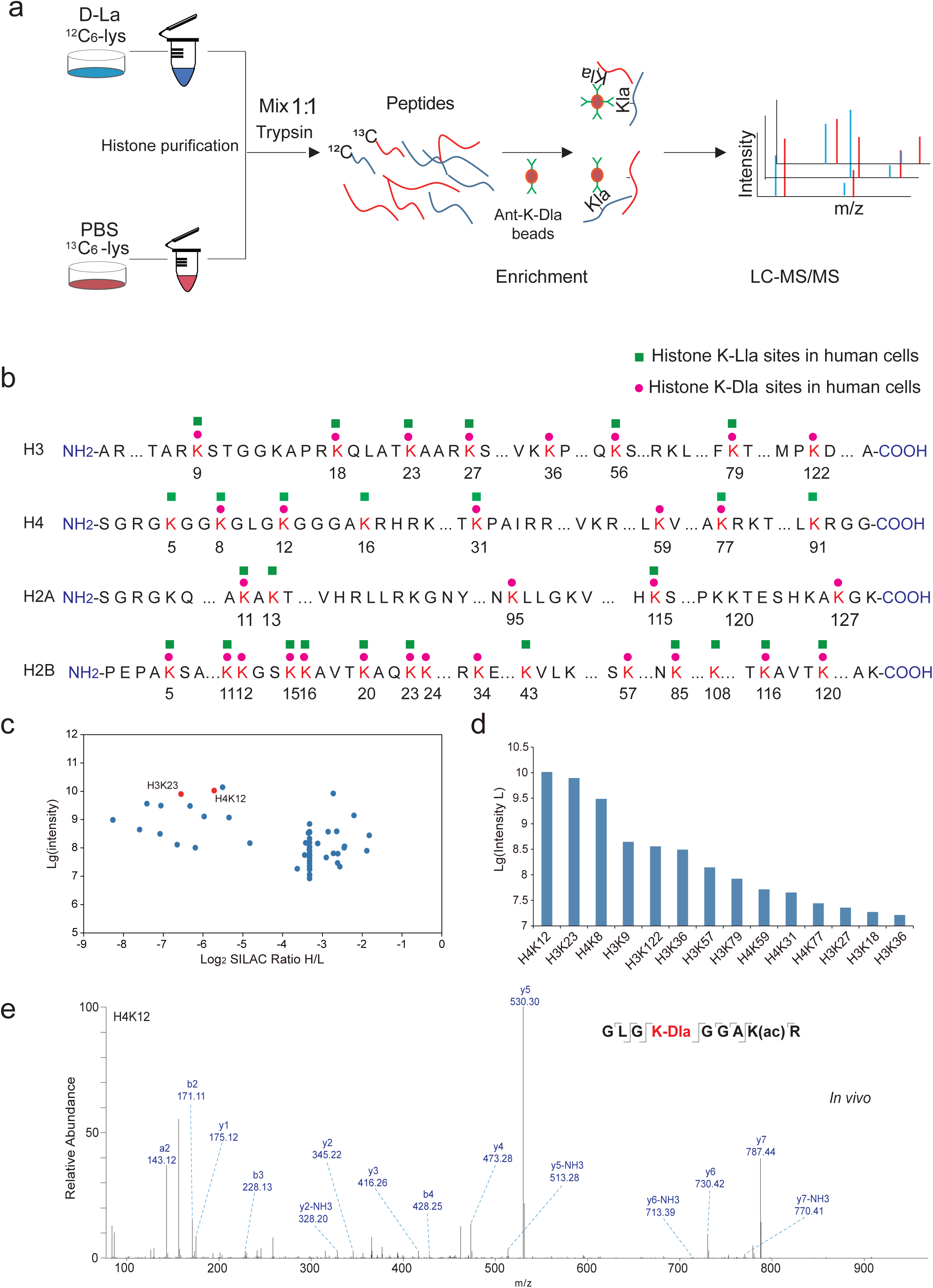
The landscape of histone D-lactylation triggered by D-la. a, The experimental workflow of SILAC quantitative proteomics to analyze the landscape of histone K-Dla regulated by D-la in HepG2 cells. b, Illustration of histone K-Dla sites identified in HepG2 cells stimulated by D-la. c, Bar graphs showing the intensity of K-Dla histones in HepG2 cells. d, Scatterplot showing the ratio and intensity of K-Dla histones in HepG2 cells. e, MS/MS spectra of a D-lactylated histone peptide (H4K12la) derived from HepG2 cells stimulated by D-la.

Meanwhile, we observed that K-Dla occurs at nearly all sites currently identified for K-Lla, suggesting that D-la-driven K-Dla is widespread on histone. To understand the preference of K-Dla driven by D-la, we calculated and compared the levels of K-Dla on histone proteins (Fig. 4c, d). We then found that K-Dla sites with the high abundance were H4K12, H3K23, H4K8, and H3K9, respectively (Fig. 4d), suggesting the D-la-driven K-Dla may be involved in the regulation of gene transcription via these histone markers. We also chemically synthesized the K-Dla peptide and compared its MS/MS fragmentation and LC retention time with its in vivo-derived counterpart (Fig. 4e and Extended Data Fig 4a-c), confirming our identifications.

### Genome-wide distribution and analysis of K-Dla substrates proteins

Our study has found that K-Dla substrates proteins were enriched in nuclear and mainly focused on histone proteins, leading us to hypothesize that K-Dla proteins can modulate gene transcription. To test the hypothesis, we performed a MNase ChIP-seq analysis by pan-anti-K-Dla antibody to identify candidate genes regulated by K-Dla substrate proteins in HepG2 cells stimulated by D-La. In contrast to control groups, we detected 19915 K-Dla binding regions in D-la stimulated HepG2 cells (Fig. 5a). Genomic distribution analysis of these K-Dla-binding regions revealed that 25.01% were located within gene promoter sequences (<=3Kb), whereas 50.4% and 19.85% were located with intronic and intergenic sequences (Fig. 5b, c).

**Fig. 5.**
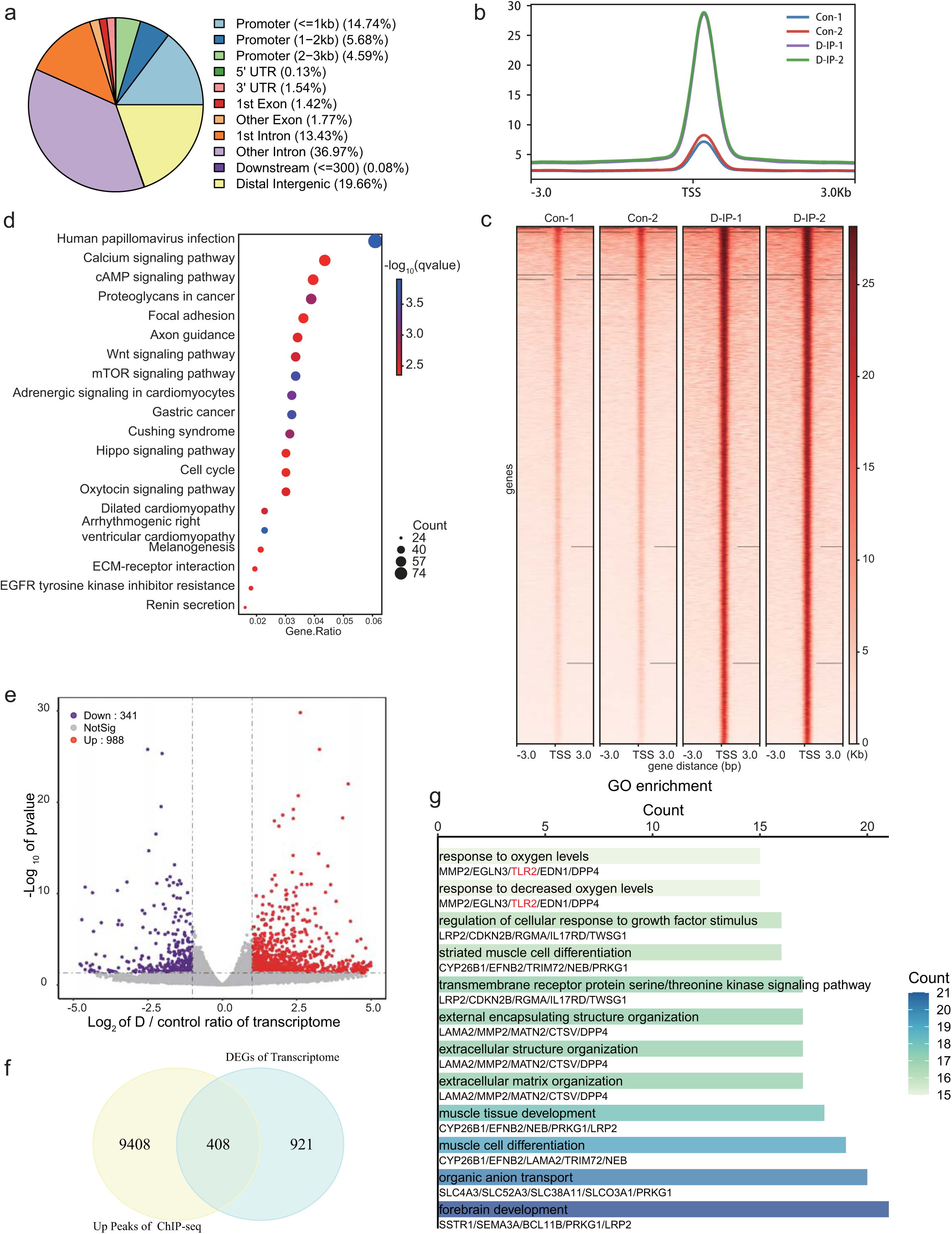
Genome-wide analysis of the transcriptional consequences of K-Dla substrates proteins. a, Genome-wide distribution of K-Dla substrates proteins binding peaks in HepG2 cells stimulated by D-la. b-c, The peaks and heatmaps presents the signal intensity which were the different K-Dla binding proteins peaks in D-IP groups. d, GO enrichment analysis of 4144 target genes of K-Dla binding regions at promoter sequences. e, Volcano map presents that 988 mRNAs were up-regulate and 341mRNA were down-regulate in D-la stimulated group (*P*<0.05, |Log 2 FC|>1). f, 408 genes were observed in an overlapped combination of DEGs and K-Dla different peaks in HepG2 cells stimulated by D-la. g, GO enrichment analysis of the overlapped genes.

To delve into the epigenetic modulatory impacts of K-Dla in HepG2 cells, a total of 4144 target genes of K-Dla binding regions at promoter sequences were enriched by gene ontology (GO) and Kyoto Encyclopedia of Genes and Genomes (KEGG) pathways. As shown in Figure 5d, Human papillomavirus infection, mTOR signaling pathway, Wnt signaling pathway and Hippo signaling pathway are enriched in KEGG pathway enrichment. Meanwhile, cell-cell signaling by wnt, Wnt signaling pathway and axonogenesis are listed in the top 3 GO terms in biological process (BP); nuclear envelope, cell leading edge and early endosome are listed in the top 3 GO terms cellular component (CC); GTPase regulator activity, nucleoside-triphosphatase regulator activity and protein serine/threonine kinase activity are in the top 3 GO terms in molecular function (MF) (Extended Data Fig. 5a).

To examine the effects of D-la in HepG2 cells, we performed gene expression analysis by RNA-seq in D-la stimulated and control cells. The results showed that 988 genes were upregulated and 341 genes were downregulated in D-la stimulated cells compared with control cells (Fig. 5e, f).

To determine whether the DEGs were induced by K-Dla in HepG2 cells, we integratively analyzed MNase ChIP-seq and RNA-seq data and revealed that 408 genes contained K-Dla binding regions that are significantly altered in D-la stimulated HepG2 cells (Fig. 5g). GO analysis showed that these genes were enriched in processes related to forebrain development, organic anion transport and response to decreased oxygen levels (Extended Data Fig. 5b).

### D-la-driven K-Dla modulates the expression of TLR2 to inhibit the HCC tumor progression

Recent study has shown that D-la can inhibit HCC tumor growth by transforming M2 tumor-associated macrophages (TAMs) to M1, but the direct function of D-la in HCC cells remains unknown.^10^ To test the effect of D-la on HCC progression, HepG2 cells were stimulated by 20 mM D-la or not, respectively. CCK-8 assay showed that D-la obviously inhibited the cell viability (Fig. 6a). Meanwhile, we found that D-la significantly reduced HepG2 cells migration and invasion (Fig. 6c and Extended Data Fig. 6a). Furthermore, the colony formation rate of HepG2 cells decreased with the increasing concentration of D-la (Fig. 6b and Extended Data Fig. 6b).

**Fig. 6.**
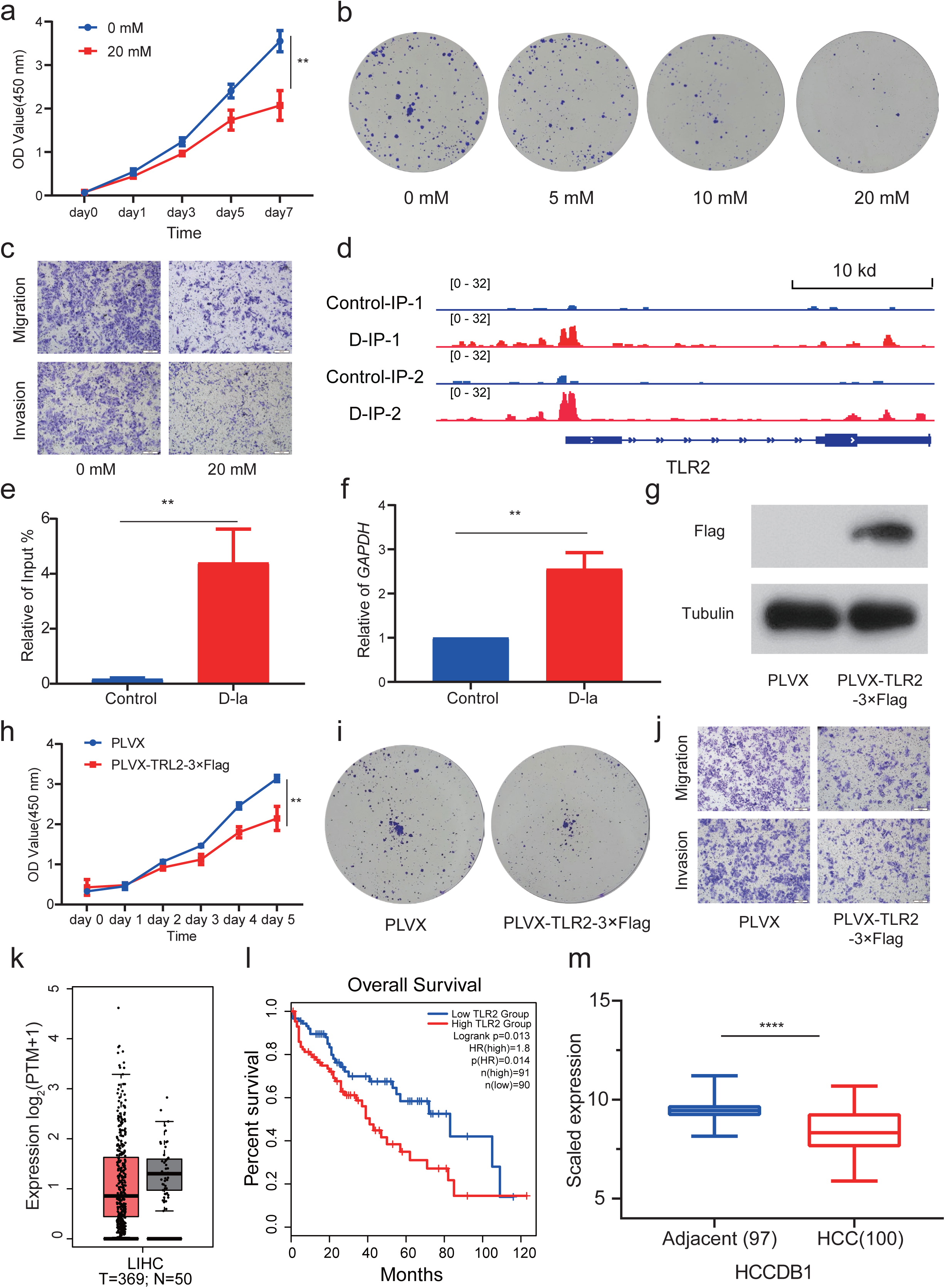
D-la-driven K-Dla modulates the expression of TLR2 in HepG2 cells to inhibit the HCC tumor progression. a, CCK-8 assay showed that D-la obviously inhibited the cell viability. b, D-la decreased the colony formation rate in HepG2 cells. c, D-la significantly inhibited HepG2 cells migration and invasion. d, TLR2 promoter in the genomic position was identified to enrich in K-Dla peaks when HepG2 cells stimulated by D-la. e, The ChIP-qPCR assays showed that K-Dla levels on TLR2 promoter were significantly elevated in HepG2 cells stimulated by D-la. f, The qPCR assays monitoring expression of the TLR2 in HepG2 cells stimulated by D-la. g, Western blotting analysis the overexpression of TLR2 in HepG2 cells. h, TCGA-LIHC data showed that TLR2 mRNA levels was lower expressed in tumors compared to the normal controls. i, TCGA-LIHC data suggested that highly expression of TRL2 influenced the Overall Survival of HCC patients. j, HCCBD database showed that TLR2 mRNA levels was lower expressed in tumors compared to the normal controls. k, CCK-8 assay showed that overexpress TLR2 in HepG2 cells inhibited the cell viability. l, Overexpressed TLR2 decreased the colony formation rate in HepG2 cells. m, Overexpressed TLR2 inhibited HepG2 cells migration and invasion.

To explore the potential mechanism of D-la inhibit HCC tumor progression, we combined the RNA-seq, MNase ChIP-seq and TCGA data base analysis. We found that the TLR2, a suppressor gene, was upregulated in RNA-seq and its promoter region enriched in K-Dla peaks (Fig. 6d). To validate the regulation of K-Dla in the expression of TLR2, we used chromatin immunoprecipitation followed by quantitative PCR (ChIP-qPCR) and qPCR to confirm our MNseq ChIP-seq and RNA-seq results. The ChIP-qPCR results showed that K-Dla level on TLR2 promoter was significantly increased in HepG2 cells stimulated by D-la (Fig. 6e). And the qPCR results showed the level of TLR2 mRNA was increased in D-LA stimulated groups too (Fig. 6f). Analysis of TCGA-LIHC data suggested that highly expression of TRL2 significantly decreased the Overall Survival of HCC patients (Fig. 6h, i). Integrative Molecular Database of Hepatocellular Carcinoma (http://lifeome.net/database/hccdb/home.html) also showed the expression of TRL2 was lower in HCC compared with adjacent (Fig. 6j and Extended Data Fig. 6c, d).

To investigate the roles of TRL2 in HCC, we overexpressed TRL2 in HepG2 cells and further performed a series of experiments for tumor behaviors (Fig. 6g). CCK-8 assay showed that overexpressed TLR2 could inhibit the cell viability (Fig. 6k). Meanwhile, we found that TLR2 reduced HepG2 cells migration and invasion (Fig. 6l and Extended Data Fig. 6e). Furthermore, overexpressed TLR2 decreased the colony formation rate of HepG2 (Fig. 6m and Extended Data Fig. 6f).

Collectively, these results showed that D-la may inhibit HCC tumor progression through a K-Dla-mediated transcription regulation for the expression of TLR2.

## Discussion

In the human body, D-la, mainly derived from intestinal bacterial metabolism, always has a lower level than L-la in normal physiological processes, because D-la might be transformed to pyruvate under LDHD catalysis in mammalian cells.^4^ Recent study also suggests that the LDHD missense variants can cause its enzymatic loss-of-function, and thus increasing D-la concentration in patient plasma.^25^ More importantly, D-la level may rapidly arise in certain human intestinal injury diseases, such as short bowel syndrome.^26^ The shortened small intestine is unable to fully absorb carbohydrates, which are then transported to the colon, where the intestinal microflora metabolizes them into large amounts of D-la in hypoxic environment, allowing to be absorbed into bloodstream.^25^ A series of neurological symptoms, such as speech disorder, gait disturbance, and changes in mental state and behavior, will happen when D-la levels in plasma are increased significantly.^25^ Some studies have shown that the concentration of D-la decreases when blood from the portal vein passes through the liver, suggesting that D-la may be stored or metabolized through the liver.^6^ Interestingly, recent report indicates that D-la acts as an endogenous immunomodulatory agent to modulate M2 tumor-associated macrophages in hepatocellular carcinoma.^10^ However, direct effect of D-la on the HCC remains unknown. Herein, we show that the secreted D-la induces and divers the generation of K-Dla for regulation of liver cancer cell behaviors, provide new insight into the function and regulatory mechanism of D-la secreted by intestinal bacteria.

Recent researches demonstrate that L-la can induce K-Lla (lysine L-lactylation) plays a key role in the transcription regulation in tumor microenvironment.^15, 24, 27, 28^ However, as the enantiomer of L-la, whether the secreted D-la can induce the K-Dla is not yet clear presently.^27, 29^ In this study, we hypothesized and verified that D-la secreted by intestinal microbiota can induce K-Dla to regulate transcription in host. Furthermore, we used four different orthogonal analytical methods to validate the existence of K-Dla in proteins. In addition to in vivo and synthetic peptides have the same chemical properties in terms of chromatographic elution in HPLC and fragmentation pattern in MS/MS, we confirmed that the K-Dla has special immunogenicity and K-Dla can be produced in cells when stimulated by D-la. Notably, we demonstrated that D-la can form K-Dla through metabolic pathways via a metabolic labelling experiment. Besides, we validate that the existence of K-Dla in liver and intestinal tissue. These results demonstrate that the secreted D-la can diver the generation of K-Dla in host.

Next, we revealed the molecular mechanism underlying K-Dla induced by exogenous D-la. We found that MCT1 is responsible to transport D-la across the plasma membrane.^21^ Inspired by our previous discovery that YdiF, a CoA transferase in *E.coli* can act as a lactyl-coenzyme,^30^ we further found that human SCOT1, holding a similar spatial structure with YdiF, can transfer coenzyme A from succinyl-CoA to D-la, and produce D-la-CoA. Surprisingly, In vitro experiments showed that SCOT1 can transfer coenzyme A from succinyl-CoA to D lactate, producing D-la-CoA. Overexpressed or knockdown SCOT1 in cells also influence the levels of K-Dla, supporting our analysis. Traditionally, SCOT1 is a key enzyme for ketone body catabolism, and can transfer coenzyme A from succinyl-CoA to acetoacetate, generating acetoacetyl-CoA. Here, we show SCOT1 has a D-lactyl-coenzyme activity, providing the key evidence for the mechanism of D-la-driven K-Dla.

Surprisingly, we revealed the K-Dla landscape with 2895 K-Dla sites on 1215 proteins. The substrates of the protein modification mainly localized to the cell nucleus and enriched in RNA related biological processes, such as mRNA progressing and RNA splicing. To characterize the proteomics of D-la-driven K-Dla, we compared it with L-lactylome in liver cancers, as well as K-Dla from S-D-(*R*)-lactoylglutathione. Thus, we found that K-Lla substrates proteins more represented in the cytosol and enriched in metabolism pathways.^24^ The unique K-Dla proteins mainly classed in RNA metabolism and gene-specific transcriptional regulator, while the unique K-Lla proteins mainly classed in metabolite interconversion enzyme and protein modifying enzyme.

Recent study has shown that methylglyoxal, which is a glycolytic by-product, can be transformed to trace amount of S-D-(*R*)-lactoylglutathione that reacts with the side chain of lysine to product the K-Dla on proteins. Although this study has not identified the precise sites of K-Dla, it predicted that the potential substrates of K-Dla from S-D-(*R*)-lactoylglutathione mainly are enriched in glycolytic enzymes by using a chemical probe.^16^

Distinguished from the above origins and pathway, our study identifies the 2895 K-Dla sites by proteomics analysis and show the D-la-driven K-Dla mainly located in nucleus (58%), especially 34 K-Dla sites on core histones are promoted by D-la. Together, the data not only suggests that D-la-driven K-Dla is a unique post-translational modification, preferentially enriched in nuclear proteins, but also provides a framework and repertoire of potential functionally D-la-targeted substrates for K-Dla.

Given that D-la-driven K-Dla tends to happen in nuclear, especially in the histones and the primary function of histone lysine PTMs is regulating target gene expressions, we therefore used MNase ChIP seq and RNA-seq to indicate the function of K-Dla in transcription regulation. Interestingly, our results showed that 19915 K-Dla binding regions were enriched in D-la stimulated HepG2 cells, with 25% located within gene promoter sequences. Integrative analysis of MNase ChIP-seq and RNA-seq data revealed that 408 genes containing K-Dla binding regions were significant changed in D-la stimulated HepG2 cells. These genes are enriched in response to oxygen levels, regulation of BMP signaling pathway, negative regulation of coagulation, and so on. Interestingly, we also found that D-la can regulate the expression of TRL2 to inhibit the HCC progression through K-Dla.

In summary, we uncover D-lactylation induced by the secreted D-la and reveal a D-la-driven K-Dla pathway, in which SCOT1 can catalyze the transform of D-la to D-lactyl-CoA, serving as a key donor for generation of K-Dla. Furthermore, we identified the K-Dla landscape with 2895 sites and characterized the K-Dla localization, enriched molecular network and pathway in cells, providing a framework and repertoire of potential functionally D-la-targeted substrates for K-Dla. Next, we analyzed the substrates differences of K-Dla and K-Lla, suggesting that D-la-driven K-Dla is a unique post-translational modification, preferentially enriched in nuclear proteins. Subsequently, we identified 34 core histone K-Dla sites and explored the effect of K-Dla on inhibiting the proliferation of hepatocellular carcinoma cells. Our study uncovered a D-la-driven K-Dla pathway and revealed a transcription regulation mediated by K-Dla for tumor suppression, providing new insight into the protein chemical modifications induced by bacterial metabolism and function of D-la secreted by gut microbiota.

## Methods

### SILAC Labeling and Cell culture

HepG2 cells were cultured in DMEM (VivaCell) medium containing 20% fetal bovine serum (FBS, VivaCell). Hela and 293T (Procell Life, CL-0005) cells were cultured with DMEM supplemented 10% FBS. All mediums supplemented with 1% antibiotics (Solarbio). All cells were cultured at 37 °C in a humidified atmosphere with 5% CO2, 21% O2 and 95% N2. HepG2 cells were grown in lysine and arginine deficient DMEM (Thermo, A33822), supplemented with 20% dialyzed FBS (Thermo), 100 U/ml of penicillin and streptomycin (Solarbio), 2 mM ^12^C6 -L-Arginine (Thermo, 89989), and 2 mM ^13^C6 -L-lysine (Thermo, 88209) or ^12^C6 -L-lysine (Thermo, 89987). Each population were grown for more than five population doublings. And the labeling efficiency was greater than 98%.

### HepG2 cells were stimulated by bacterial medium or ^13^C1-D-lactate

The DMEM cell medium containing 20% serum was mixed 1:1 with the filtrated bacterial solution supernatant. The mixture was added to HepG2 cells. The cells were cultured for 48 hours in 37°C incubator.

Treatment of ^13^C1-D-lactate. To verify whether D-la is the precursor of histone K-Dla, ^13^C1-D-lactate (Sigma, 676144) was added to HepG2 cells. Cells were treated with 20 mM ^13^C1-D-lactate for 48 hours before collecting proteins.

### Detection of L/D-lactate for the supernatant of *E. coli* culture medium

Firstly, *E. coli* strain MG1655 (Tolo Biotech) was cultured overnight in LB medium (Solarbio). Then, 200 ul *E. coli* culture medium was inoculated into anaerobic and aerobic culture tubes with M9 (Sangon) medium, respectively, and cultured overnight at 37°C. After cultivation, centrifuge 8000 g at 4°C for 10 minutes, collect the supernatant. Subsequent experiments were conducted to detect the concentrations using L/D-lactate colorimetric assay kits (Elabscience).

### Clearance of gut microbiota in mice

Mouse protocols were approved by the Laboratory Animal Management and Use Committee (IACUC) of Tianjin Medical University. The 6-8 weeks old male C57BL/6 mice were maintained at 25°C and on an artificial 12-hlight–dark cycle. Mice were randomly assigned to different groups. Mice drank waters and ate foods ad libitum.

For the gut microbiota depleting assay, mice were treated with broad-spectrum antibiotics (ATBs, a mixture of metronidazole (1 g/L), neomycin (1 g/L), ampicillin (1 g/L), and vancomycin (0.5 g/L)) in drinking water ad libitum and (a mixture of metronidazole (200 mg/kg), neomycin (200 mg/kg), ampicillin (200 mg/kg), and vancomycin (100 mg/kg)) via oral gavage once a day for 7 days. The DNA in the mouse feces was obtained using the fecal DNA extraction kit for the PCR experiment to test the sterilization effect (27F:AGAGTTTGATCCTGGCTCAG, 1492R: GGTTACCTTGTTACGACTT).

### RNA Sequencing and Real-Time Quantitative PCR

HepG2 cells were cultured in DMEMs stimulated by 20 mM D-la for 48 hours. Total RNA was isolated from cells using TRIzol reagent (Thermo). cDNA was generated using StarScrip (Genstar,) and qPCR was performed using RealStar Fast SYBR qPCR Mix (Genstar,). RNA-seq was performed on the Illumina NovaSeq platform. Differential expressed genes were calculated by DESeq2. The *P* was less than 0.05 and the absolute of fold change more than 2 were considered as significant differential expression genes. GO and KEGG Pathway analysis were performed for differentially expressed genes. The relative expression of target genes were normalized to GAPDH via the 2^-△△ct^ method.

### Whole-cell protein and histone extraction

Cells were lysed by RIPA (Beyotime) supplemented with Protease Inhibitor Cocktail (Roche), PMSF (Sangon) and deacetylase inhibitor (Beyotime). Protein concentration was quantified by bicinchoninic acid assay (Thermo Fisher).

Histones from HepG2 (Pricella, CL-0103) cells were extracted using an acid-extraction protocol. Briefly, cells were collected in lysis buffer (PBS with 1%NP-40, Deacetylase Inhibitor Cocktail, 1 Mm Protease Inhibitor Cocktail and 1 mM PMSF) and revolve slowly for 15 min. The nuclei were collected by centrifugation at 1000 g for 15 min at 4 °C and wash the nuclei by lysis buffer, centrifuge, and discard the supernatant. Add 0.2 mol/L H2SO4 solution of about 500 μl to dissolve the histone protein and gentle rotation over night at 4°C. The mix were centrifuged at 1,6000 g for 15 min at 4°C.The supernatants were collected and mixed with 25% (final concentration) trichloroacetic acid (TCA, Aladdin) on ice. The mix stand for 30 min on ice. The mix was centrifuged at 1,6000 g for 30 min at 4°C. After acetone washing thrice and drying, the histone was dissolved by 100 mM NH4HCO3 for next experiment.

### Western blot assay

Western blot analysis was performed as previously described. Samples were denatured in SDS loading buffer and heated at 98°C for 10 minutes. Protein samples were separated via 12% or 15% SDS-PAGE. Electrophoresis was used to isolate proteins. Then, it is transferred to nitrocellulose filter membrane, which was blocked with 5% skim milk at room temperature for 2 hours. Antibodies were incubated with membranes overnight at 4°C as described: K-Dla (PTM, PTM1429D), K-Lla (PTM, PTM-1401RM), SCOT1 (proteintech, 12175-1-AP), Flag (proteintech, 66008-4-Ig), α-Tubulin (proteintech, 11224-1-AP), Kac (PTM, 200-101-232S). Following 3× washes with TBST (Genestar) and membranes incubated with horseradase-labeled secondary antibodies (ZSGB-bio ZB-2306 or ZB-5305, 1:10,000) for 2 hours at room temperature. Following 3x washes with TBST. Using the fluorescence EDL enhanced chemical luminant (Genestar) to detect the proteins and using the X-films to image the bands.

### The enrichment of K-Dla of whole-cell proteins

As described previously (Extended Data Fig. 2A), HepG2 cells were cultured in DMEMs stimulated by 20 mM D-la for 48 hours. Cells were collected and the whole proteins of the cells was extracted. The appropriate of proteins are precipitated by 25% (final concentration) TCA for 2h on ice. The mix was centrifuged at 1,6000 g for 30 min at 4°C. After acetone washing thrice and drying, the precipitated proteins were dissolved by 100 mM NH4HCO3 and digested overnight by trypsin (trypsin: protein ratio,1: 50, Promega) at 37°C. The digestive products were added to the final concentration of 5 mM DTT (Sigma) and incubated for 1 h at 56°C, 15 mM iodoacetamide was incubated under darkness at room temperature for 45 min, and then 30 mM cysteine (Sangon) was incubated at room temperature for 30 min to block excess iodoacetamide (Thermo). Then secondary digestion was performed with trypsin (1: 100) at 37°C for 4 h. The product was desalted by SepPak C18 cartridges (Strata-X) and subsequently drained by the Speed Vac system (Thermo). Immunoaffinity enrichment of lysine D-lactylation was then performed.

### The enrichment of K-Dla of histone in SILAC samples

As described in Figure 2A, ^12^C6 -L-lysine labeling HepG2 cells were cultured in DMEMs stimulated by 20 mM D-la for 48 hours and ^13^C6 -L-lysine labeling HepG2 cells were cultured in DMEMs stimulated by PBS. Cells were harvested to extract histones. Two group histones mixed 1:1 and digested overnight by trypsin (trypsin: protein ratio, 1:50). 10% TFA terminate digestion and products were desalted by SepPak C18 cartridges and dried. Immunoaffinity enrichment of lysine D-lactylation was then performed.

### Immunoaffinity enrichment

Immunoaffinity enrichment was performed as described previously.^30^ In detail, the drained samples were redissolved in NETN buffer (50 mM Tris-HCl pH 8.0, 100 mM NaCl, 1 mM EDTA, 0.5% Nonidet P-40) and incubated with pre-washed anti-D-lactyllysine antibody-conjugated protein A agarose beads (PTM Biolabs) at 4 °C overnight, with gentle rotation. The incubated beads were washed three times with NETN buffer, twice with ETN buffer (50 mM Tris-HCl pH 8.0, 100 mM NaCl and 1 mM EDTA) and three times with water. The enriched peptides were eluted three times with 1% trifluoroacetic acid. Finally, the eluates were drained and desalted with C18 ZipTips (Millipore Corp) before nano-HPLC–MS/MS analysis.

### HPLC-MS/MS analysis for K-Dla

The enriched peptides were analyzed by HPLC-MS/MS. Samples was reconstituted in 0.1% formic acid and then injected into a nano-LC system (EASY-nLC 1200, Thermo Fisher Scientific) using trap-elute mode. After loading into the trap column (Thermo Scientific Acclaim PepMap 100 C18, 75μm*2cm, 3μm, 100Å), all of the peptides were further separated by a home-packed 75 µm-i.d., 25 cm-long C18 (1.9 µm, Dr. Maisch GmbH, Ammerbuch, Germany) column at flow rate of 300 nL/min. Gradient elution was performed with 5-7% HPLC buffer B (0.1% formic acid in 80% acetonitrile) for 2 min, 7%-22% buffer B for 78 min, 22%-40% buffer B for 38 min, 40%-100% buffer B for 3min and keep with 100% buffer B for 9 min. Spray voltage was set to 2.1 kV, funnel RF level at 40, and ion transfer tube temperature at 320 °C. Mass spectrometric analysis was carried out in data-dependent acquisition (DDA) mode of the most intense precursors, and data were collected by Xcalibur (v.4.0.27.19). The orbitrap mass analyzer was used as the MS1 detector with 60,000 resolution and scan range 350–1750 m/z. The normalized AGC target and maximum injection time were set at 100% /50 ms for MS1, and 200% /50 ms for MS2. The orbitrap mass analyzer was used as the MS2 detector with 17,500 resolutions.

Precursor ions with charges of +2 to +5 were isolated for MS2, and dynamic exclusion time was set at 50 s. The MS2 isolation window was 1.6 Da, and a normalized HCD (higher-energy collision-induced dissociation) collision energy of 30% was used for precursor fragmentation.

### Database search and data filter criteria for K-Dla

The database search and filter criteria were performed as described previously.^30^ In detail, raw data were searched by MaxQuant (v.1.5.5.1) with UniProt Homo sapiens protein database (Proteome ID: UP000005640) and an overall false discovery rate for peptides of less than 1%. Peptide sequences searching was set as trypsin specificity and six for minimal peptide length. Two missed cleavages for maximum were set for whole cell proteins K-D-la, and five missed cleavages were set for the histones K-D-la. Carbamidomethylation on Cys was specified as fixed modification. D-lactylation on lysine, oxidation of methionine and acetylation on the protein N-terminus were set as variable modifications. Mass tolerances were set at ±10 ppm for precursor ions and ±0.02 Da for MS/MS. D-lactylated peptides with a score of <40 and localization probability of <0.75 were further excluded.

### Purification of recombinant proteins

Transformed pET28a-SCOT1 vectors into *E.coli BL21* (Sangon).The *E.coli BL21* were cultured in LB medium containing kanamycin (Kana, 50 μg/ml), at 37 in shaken flasks to an optical density of 0.6-0.8 at 600 nm. Next, cells were induced with 0.1 mM IPTG at 16°C overnight. Then harvested the whole-cell lysates for protein purification. The cells were split by lysis buffer (20 mM Tris-HCl, pH 8.0, 10 mM MgCl2, 1 mg/ml lysozyme, 50 U/ml nuclease) and ultrasonication for 30 min (power 25%, 30 s on and 30 s off). The supernatants were collected by centrifugation at 12000 g for 5 min at 4 °C. Then the supernatants were mixed with HisPur Ni-NTA Resin and washed by wash buffer (20 mM Na3PO4, 300 mM NaCl, 25 mM imidazole, pH 7.4), the recombinant proteins were eluted with elution buffer (20 mM Na3PO4, 300 mM NaCl, 250 mM imidazole (Sangon), PH 7.4). The elution was collected and concentrated using an Amicon Ultra-0.5 Centrifugal Filter Device (Sigma) in storage buffer (15 mM MgCl2, 50 mM Tris-HCl, PH 9.1).

### In vitro assay for SCOT1 activity

Assay of the SCOT1 CoA-transferase activity was performed as described previously.^30^ In detail, the enzyme reactions were carried out in SCOT1 buffer [15 mM MgCl2, 50 mM Tris-HCl, PH 9.1] containing 1 mM Succinyl-CoA and 10 mM D/L-lactate at 35 °C for 20 min, and initialized by adding the 7.5 μg of purified OXCT1. The total volume of the reaction mixture was 50 μL. The reactions were terminated by adding equal volume of 10% (v/v) trifluoroacetic acid and centrifugation, the supernatant was cleaned with C18 ZipTips, and drained by the Speed Vac system (Thermo Fisher Scientific). The samples were analyzed by LC-MS/MS. Each reaction was performed three biological repetitions

### UPLC-MS/MS analysis for D-lactyl-CoA

The drained samples dissolved in water were analyzed by an Orbitrap Exploris 480 (Thermo Scientific) mass spectrometer. In detail, samples were separated by a ACQUITY UPLC BEH C18 column (2.1*100 mm,1.7 μm), the HPLC gradient for 15 min was set up as follows: 0.2 mL/min flow at 98% buffer A (water with 5 mM ammonium acetate) and 2% buffer B (95 % acetonitrile in water with 5 mM ammonium acetate) for 0 min, 98 to 70% buffer A for 8 min, 70 to 2% buffer A for 1 min, 2% buffer A for 3 min, 2 to 98% buffer A for 1 min, 98% buffer A for 2 min. An Orbitrap Exploris 480 (Thermo Fisher Scientific) was employed for MS analysis in positive ESI mode. Spray voltage was set to 3.5 kV, funnel RF level at 40, and ion transfer tube temperature at 320 °C. Data were collected by Xcalibur (v.4.0.27.19). The orbitrap mass analyzer was used as the MS1 detector with 120,000 resolution. The normalized AGC target and maximum injection time were set at 70% /standard for MS1. The MS2 detector was used 15,000 resolution. Targeted mass setting is D-lactyl-CoA: 840.1436. MS2 isolation window was 2 Da and mass tolerance was 10 ppm, and a normalized HCD (higher-energy collision-induced dissociation) collision energy of 25 % was used for precursor fragmentation. The relative intensity of D-lactyl-CoA was analyzed by Xcalibur (v.4.0.27.19)

### MNase ChIP-seq

The MNase ChIP-seq was performed using a gold-standard’ ChIP protocol.^2, 27, 28^ In brief, 5X10^6^ cells were fixed with 1% formaldehyde at room temperature and revolve slowly for 10 minutes. Crosslinking stopped by 150 mM of glycine for 5 minutes at room temperature. Cells were washed twice with cold PBS and then the nuclei were collected by centrifugation at 700 g for 3 min at 4 °C. Chromatin was digested in 1 μl MNase (Sigma) at 37 °C for 20 min. Reaction stopped by 0.5 M EDTA. Immunoprecipitation was performed using 1 μl of K-Dla antibody with protein A/G beads (Solarbio) gentle rotation overnight at 4°C. After elution and reversed crosslinking, samples were treated with RNase A (Solarbio) for 30 minutes at 37°C and Proteinase K (Sangon) for 2 h at 55 °C. DNA was purified by phenol chloroform and 5 ng of raw ChIP material was processed for library construction. For ChIP sequencing (ChIP-seq), sequenced on the Illumina NovaSeq platform.

### ChIP-seq analysis

To assess the quality of ChIP-seq data, FastQC tool was used to examine metrics, such as sequencing error rate, sequencing depth, and fragment length distribution. Preprocessing steps on the raw sequencing data including removing low-quality sequences and adapter trimming were performed. Aligned the preprocessed reads to a reference genome using alignment tool (Bowtie2). Peak calling was performed using the model-based analysis of ChIP–seq (MACS2). To identify sites with differential enrichment between the two experimental groups, we utilized the R package DiffBind. The mapping results were visualized using the UCSC Genome Browser on Human GRCh38/hg38. The annotation of ChIP-seq peaks was performed using the ChipSeeker package.

### Immunofluorescence staining

Cells were washed by cold PBS and fixed using 4% paraformaldehyde (Solarbio) for 15 min at room temperature. The samples were washed 3 times with cold PBS and permeabilized with 0.25% Triton X-100 (Solarbio) for 15 min at room temperature and blocked with 10% BSA (Sigma) for 2h.

Then, cells were blocked with 10% BSA or 2h. Antibodies were incubated with membranes overnight at 4°C. Following 3x washes with PBS and cells incubated with Fluorescence secondary antibody (Invitrogen, A-11034) for 1 hours at room temperature. Following 3x washes with PBS. All samples were incubated with antifade mounting medium for fluorescence including DAPI (ZSGB). The fluorescence was observed under fluorescence microscope.

### Cell clonal formation assays

Cells were counted and 1000 cells/well were inoculated into the 12-well plate and the medium was changed every 3 days. About a week after cell culture, the cells were washed twice with PBS and fixed with a mixture of methanol: acetic acid ratio 3:1. The cells were stained with 0.2% crystal violet solution (Solarbio) at room temperature for 5 mins, and the excess crystal violet solution was washed away with PBS. The formation of cell clones was photographed and counted.

### Transwell migration and invasion experiment

To perform the transwell (Corning) invasion assay, matrigel Matrix (Corning) was seeded in the upper chamber with a pore size of 8.0 μm, and the transwell placed in 37 °C until the Matrigel solidified. Then, 50,000 or 80,000cells were mixed in 200 μL serum-free medium and seeded in the upper chamber. Finally, 600 μL of 20% FBS DMEM was added to the bottom chambers. After incubation for 36 hours, cells were fixed with a mixture of methanol: acetic acid ratio 3:1 for 5 min and 0.2% crystal violet staining for10 min. After washing away the excess crystal violet solution and drying, the cells were photographed under a microscope and counted.

### CRISPR-Cas9 establishment of SCOT1 knockout Hela cells

SCOT1-sg-for: CGATCCTCCTTCTTGTACCA and SCOT1-sg-rev: TGGTACAAGAAGGAGGATCG were inserted into the plasmid to target SCOT1.The target sgRNA was cloned into the lenti-CRISPR v2 vector and co-transfected with psPAX2 and pMD2G plasmids to generate the corresponding virus in HEK293T cells by PEI. Virus was harvested after 48 h. The Hela cells were placed in a 12-well plate, and after the cells were attached to the wall, 100 μl disease venom was added into the medium. After 48 hours of virus infection, the cells were changed into fluid and added 2 μg/μl purinomycin (Solarbio) until no cells died.

### Establishment of target gene expression cells

The target gene ligated into the PLVX-Puroplasmid with an N-terminal 3×flag-tag by T4 ligase (Genstar). The variants were cotransfected with plasmids encoding PML, TAT, VSVG and REV into HEK293T packaging cells using PEI (Yeasen). Those cells were placed in a 12-well plate, and after the cells were attached to the wall, 100μl disease venom was added into the medium. After 48 hours of virus infection, the cells were changed into fluid and added 2 μg/μl purinomycin until no cells died.

## Data availability

The data supporting the findings of this study are available within the article and its supplemental information. The mass spectrometry proteomics data have been deposited to the ProteomeXchange Consortium (http://proteomecentral.proteomexchange.org) via the iProX partner repository with the dataset identifier (PXD053161). The MNase ChIP-seq and RNA-seq data generated in this study have been deposited in the Gene Expression Omnibus (GEO) repository under accession code (GSE269963).

## Code availability

The software and code used to analyze this data is all available online and has been detailed in the Methods section.

## Acknowledgements

This work was supported by the Funding of National Natural Science Foundation of China to K.Z. (22074103, 22274114), C.L. (32200735), G.Z. (22374106) and X.B. (22004091) and the Talent Excellence Program from Tianjin Medical University to K.Z.

## Author contributions

K.Z. supervised experiments, Y.Z., J.Zs and K.Z. designed experiments. Y.Z., J.Z. and K.Z. wrote the manuscript. Y.Z., J.Z., and M.X. carried out cell culture, enzymatic activity assay, and molecular biological experiments. J.Z., Y.Z., M.X. Y.H. H.Z., A.W., and K.Z. carried out proteomic survey. Y.Z., J.Z., M.X., A.W., Z.F., J.W., S.W., Z.N., Y.H., H.Z., X.B., S.T., C.C., G.Z., H.D., D.H., Z.C., and K.Z. carried out data collection, analysis and interpretation. All of the authors discussed the results and commented on the manuscript.

---

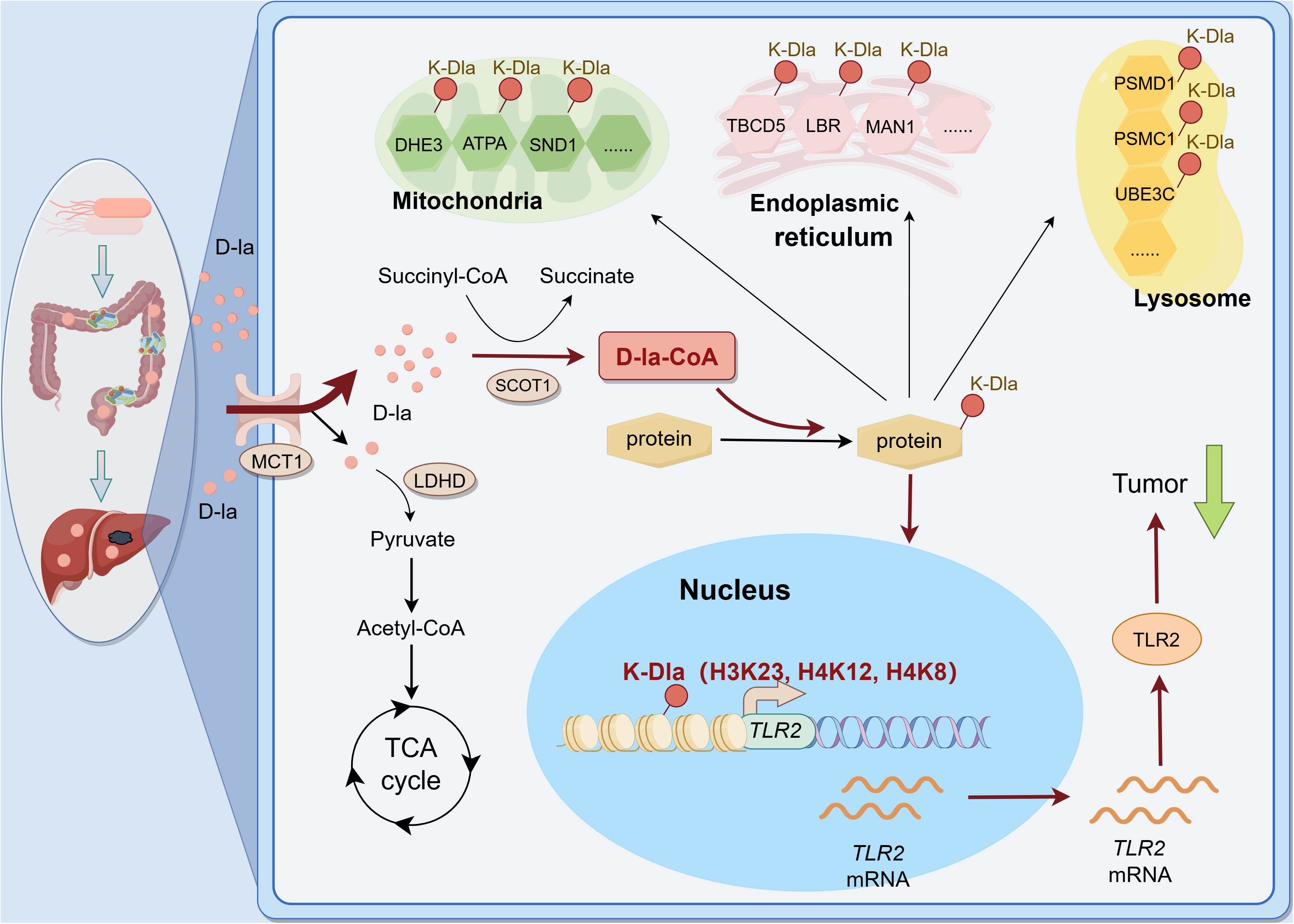

